# Ocular vestibular evoked myogenic potential (VEMP) reveals subcortical HTLV-1-associated neurological disease

**DOI:** 10.1101/634709

**Authors:** Tatiana Rocha Silva, Marco Aurélio Rocha Santos, Luciana Macedo de Resende, Ludimila Labanca, Rafael Teixeira Scoralick Dias, Denise Utsch Gonçalves

## Abstract

**Introduction:** Vestibular Myogenic Evoked Potential (VEMP) evaluates vestibulo-ocular and vestibulospinal reflexes associated with posture.

**Purpose:** To compare cervical and ocular VEMP in individuals with HTLV-1 associated myelopathy (HAM) and with HTLV-1-asymptomatic infection.

**Materials and Methods:** This cross-sectional study included 52 HTLV-1-infected individuals (26 HAM and 26 asymptomatic carriers) and 26 negative controls. The groups were similar regarding age and gender. Participants underwent ocular and cervical VEMP that were performed simultaneously. The stimulus used to generate VEMP was a sound, low-frequency toneburst, intensity of 120 decibels normalized hearing level (dB nHL), bandpass filter from 10 to 1,500 Hz, with 100 stimuli at 500 Hertz (Hz) and 50 milliseconds (ms) recording time. An alteration in the electrophysiological waves P13 and N23 for cervical VEMP and N10 and P15 waves for ocular VEMP was compared between groups.

**Results:** Cervical VEMP was different among the groups for P13 (p=0.001) and N23 (p=0.003). Ocular VEMP was similar for N10 (p=0.375) and different for P15 (p=0.000). In the HTLV-1-asymptomatic group, 1(3.8%) individual presented changes in both ocular and cervical VEMP, while in HAM group, 16(61.5%) presented changes in both tests.

**Conclusion:** Neurological impairment in HAM was not restricted to the spinal cord. The mesencephalic and thalamic connections, tested by ocular VEMP, were also altered. Damage of the oculomotor system, responsible for eye stabilization during head and body movements, may explain why dizziness is such a frequent complaint in HAM.

**Authors’ summary:** Human T-cell lymphotropic virus type 1 (HTLV-1) infection is endemic in Brazil and can cause HTLV-1-associated myelopathy (HAM). This neurological disease progresses slowly and, within ten years after its onset, can confine the patient to a wheelchair. Changes in HAM inflammatory characteristics can subsequently occur in the cortex, subcortical white matter, cerebellum, and brainstem. In the present study, we used the electrophysiological test Vestibular Myogenic Evoked Potential (VEMP) to evaluate the thalamic, brainstem, and spinal neural connections. This test evaluates the peripheral and the central vestibular pathway and has been used to test the postural reflexes involved in the control of one’s balance. The VEMP from the oculomotor muscles demonstrated that a subcortical impairment occurs in HAM and can also occur in the asymptomatic phase of HTLV-1 infection.

## Introduction

The Human T-cell lymphotropic virus type 1 (HTLV-1) infection affects approximately 5-10 million people worldwide [1]. The majority of the infected individuals remain asymptomatic throughout their lives [2]. The host genetic and immunological factors seem to be related to the development of HTLV-1-associated diseases [1,2].

The range of neurological manifestations of HTLV-1-associated myelopathy (HAM) includes not only the spine, with the classical motor limitations affecting the lower limbs, but also the autonomic dysfunction [3]. In fact, inflammatory alterations due to HAM can be detected in the cortex, subcortical white matter, cerebellum, and brainstem, mainly in the advanced phases of this disease [4–7].

The complaint of dizziness has proven to be frequent HAM and can be one of the first symptoms of HTLV-1-neurological impairment [11,12]. Therefore, individuals infected with HTLV-1 may present vague complaints, with no motor, sensitive, or autonomic abnormalities [4–6]. HAM diagnosis is based on clinical criteria that reveals established neurological damage [13].

Vestibular Evoked Myogenic Potential (VEMP) evaluates the integration of the vestibular nerves with the brainstem and the muscular system. It tests the peripheral and central vestibular pathway and has been used to test brainstem functions [14,15] and postural reflexes [12,16]. VEMP tests the vestibulospinal and vestibulo-ocular reflexes involved in the control of the postural balance. Normal VEMP depends on the functional integrity of the saccular and utricular maculae, the inferior vestibular nerve, the superior vestibular nerve, the vestibular nuclei, the central vestibular pathways, and the neuromuscular plaques involved in these reflexes [17,18].

The subclinical spinal cord injury related to HAM has been already shown through VEMP of cervical and of lower limbs muscles, exams that are used to test the vestibulospinal reflex [11,12,16,19,20]. The present study proposes the use of VEMP of the oculomotor system (ocular VEMP) to test the subcortical pathways associated with body balance to verify the extension of the HTLV-1-neurological damage.

## Methods

### Study design

The study was a comparative cross-sectional analysis. Cervical VEMP and ocular VEMP were compared between individuals with definite HAM and HTLV-1-asymptomatic carriers and healthy controls.

### Ethical aspects

This research was conducted in accordance with the principles expressed in the Declaration of Helsinki and was approved by the Research Ethics Committee from Universidade Federal de Minas Gerais (COEP UFMG), logged under protocol number CAAE 92928518.3.0000.5149. All participants provided voluntary written consent and declared that they were aware of the study procedures and their choice to participate.

### Sample size

The sample size was calculated using G* Power software 3.1.9.2 (Heinrich-Heine Universitat Düsseldorf, Düsseldorf, Germany, 2007) to achieve a power of 80% and a significance level of 5% based on the mean and standard deviation of the P13-N23 response of patients with HAM and healthy controls [11]. The final calculation included 26 participants per group.

### Participants

The groups of study were recruited from a cohort of former blood donors infected with HTLV-1 who have received follow-up from the Interdisciplinary HTLV Research Group (GIPH) since 1997, in Belo Horizonte, Brazil [21]. The GIPH evaluates the natural history, clinical manifestations and epidemiological aspects of HTLV infection.

Seventy-eight individuals, 32 to 60 years of age, were invited to participate in this study. The participants consisted of 26 individuals with definite HAM, 26 with HTLV-1-asymptomatic infection, and a control group of 26 individuals not infected by HTLV-1.

The classification of the participants infected by HTLV-1 regarding neurological impairment followed the Expanded Disability Status Scale (EDSS) [22] and the OSAME scale [23]: asymptomatic individual, (EDSS and OSAME - 0 on both scales) and definite diagnosis of HAM (EDSS and OSAME greater than 2 on both scales).

Individuals with positive serology for the Human Immunodeficiency Virus (HIV), HTLV-2, or any other blood-tested disease were excluded, as well as an undetermined serology for HTLV-1 and a positive *Venereal Disease Research Laboratory* (VDRL) test.

The control group consisted of active blood donors followed by GIPH as the negative controls. Concerning all the participants, individuals with neurological diseases, otitis, tympanic membrane perforation, history of otologic surgery or peripheral vestibular disease, as well as individuals unable to perform cervical rotation and ocular movement were excluded.

### Vestibular Evoked Myogenic Potential (VEMP)

VEMP was performed with Labat^®^ equipment, using two channels. The stimuli were presented through ER 3A insertion phones, with disposable foam eartips. Tone burst stimuli at an intensity of 120 decibels normalized hearing level (dB nHL) were used. In this study, a bandpass filter of 10 to 1,500 Hertz (Hz) was used. To obtain each record, 100 stimuli were presented at a frequency of 500 Hz at a rate of four stimuli per second. The scan window was 50 milliseconds (ms). Each subject underwent at least two stimulations per side, to verify the replication of the potential. The impedance values, which had to be below 5 kiloohm (KΩ), were checked before each record [14].

The recording of cervical VEMP and ocular VEMP was performed simultaneously. Channel 1 electrodes were used to record ocular VEMP and channel 2 electrodes to record cervical VEMP [14].

The active electrode related to cervical VEMP was placed on the opposite side at the anterior border of the sternocleidomastoid muscle in its upper third, and the reference electrode was placed in the sternal notch region. For ocular VEMP recording, the active electrode (negative electrode) in channel 1 was placed approximately 1 centimeter (cm) below the lower eyelid, and the reference electrode (positive electrode) was placed at a distance of approximately 1 cm from the active electrode. The ground electrode was placed on the forehead (Fpz) (Fig 1).

**Fig 1.**
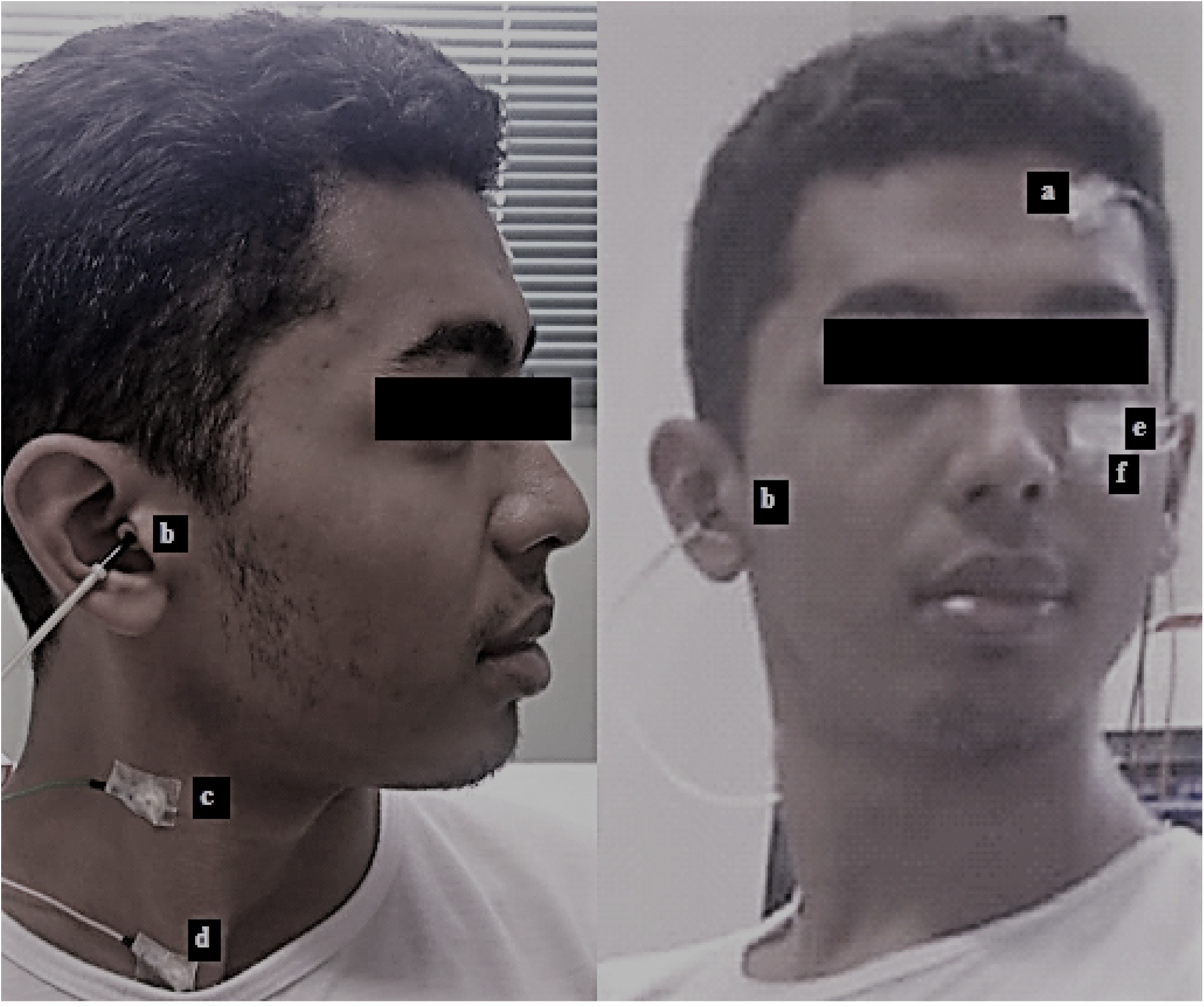
Simultaneous cervical and ocular VEMP. (a) ground electrode. (b) auditory stimulus. (c) active electrode on channel 2 at the anterior border of the sternocleidomastoid muscle in its upper third. (d) reference electrode on channel 2 at the sternal notch region. (e) active electrode on channel 1 below the lower eyelid. (f) reference electrode on channel 1 below the active electrode.

Participants were instructed to sit on the chair and keep their heads rotated to the opposite side of the stimulated ear, causing contraction of the sternocleidomastoid muscle. At the same time, the participant was instructed to look at a stationary target located on the wall in front of him and then immediately at a fixed point located above the target, which formed a vertical viewing angle of approximately 30º above the horizontal plane. The Simultaneous ocular and cervical VEMP protocol is available at dx.doi.org/10.17504/protocols.io.zmzf476.

The ocular VEMP is composed of two sets of biphasic waveforms. The first biphasic potential has a negative peak (N) with an average latency of 10 ms, followed by a positive peak (P) with an average latency of 15 ms, which is known as N10–P15. The cervical VEMP consists of two sets of biphasic waveforms. The first biphasic potential has a positive peak (P) with an average latency of 13 milliseconds (ms), followed by a negative peak (N) with an average latency of 23 ms, which it known as P13–N23 (Fig 2).

**Fig 2.**
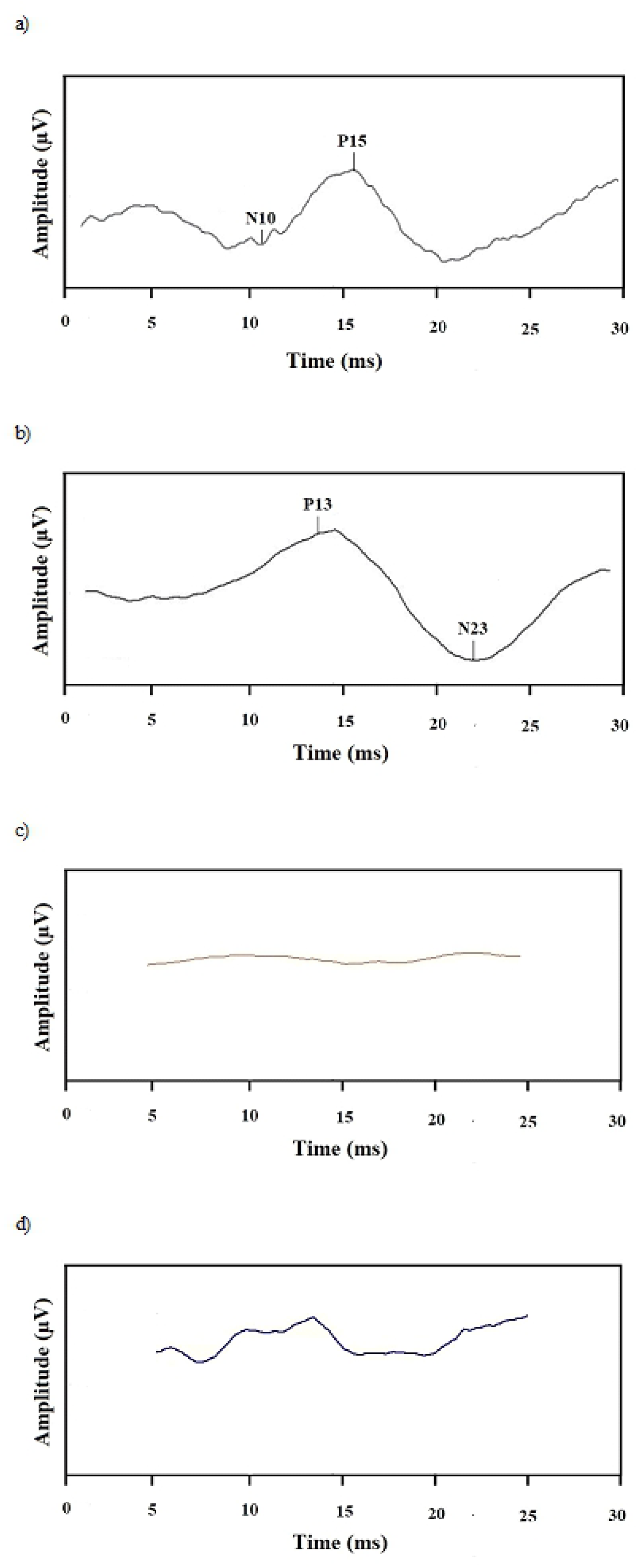
Examples of tracings obtained by the VEMP records. a) normal ocular VEMP. b) normal cervical VEMP. c) altered ocular VEMP (no response). d) altered cervical VEMP (no response).

The American Society of Encephalography and Evoked Potentials’ criteria for evoked potentials were considered for the analysis of the latency values of the cervical VEMP and ocular VEMP waves. The definition of altered latency values includes those that exceed 2.5 standard deviations (SD) [24]. In this study, for cervical VEMP, we considered the normal latency of 13 ms (±2.5 SD) for P13 and 23 ms (±2.5 SD) for N23, while for ocular VEMP, we considered the normal latency of 10ms (±2.5 SD) for N10 and 15ms (±2.5 SD) for P15 [25,26]. The validation of the analyzed reference values was guaranteed by comparing these with parameters already established in other national and international peer reviews [14,27,28].

The parameters considered in the VEMP analysis are the latency and amplitude of the waves. However, the amplitude may vary according to age, muscular strength [27,29], and cochlear diseases [30,31]. Therefore, amplitude was not considered in the analysis since this variable is not consistent to define neural conduction abnormalities.

### Statistical analysis of data

VEMP results were classified as normal and altered. Latency prolongation and no response were considered as the altered results. Ocular and cervical VEMP were compared between the groups infected and not infected by HTLV-1.

Statistical analysis was performed using the *Statistical Package for Social Sciences* (SPSS), version 20.0. The normality of the samples was assessed using the Kolmogorov-Smirnov and Shapiro-Wilk tests. The comparison between groups was performed using the Kruskal-Wallis test, Chi-square or Fisher’s Exact test, and Kruskal-wallis with Bonferroni correction. The adopted level of significance was 5% (p≤0.05).

## Results

The characteristics of the studied population and the classification in the neurological scales are described in Table 1. The groups were similar regarding gender and age.

**Table 1.**
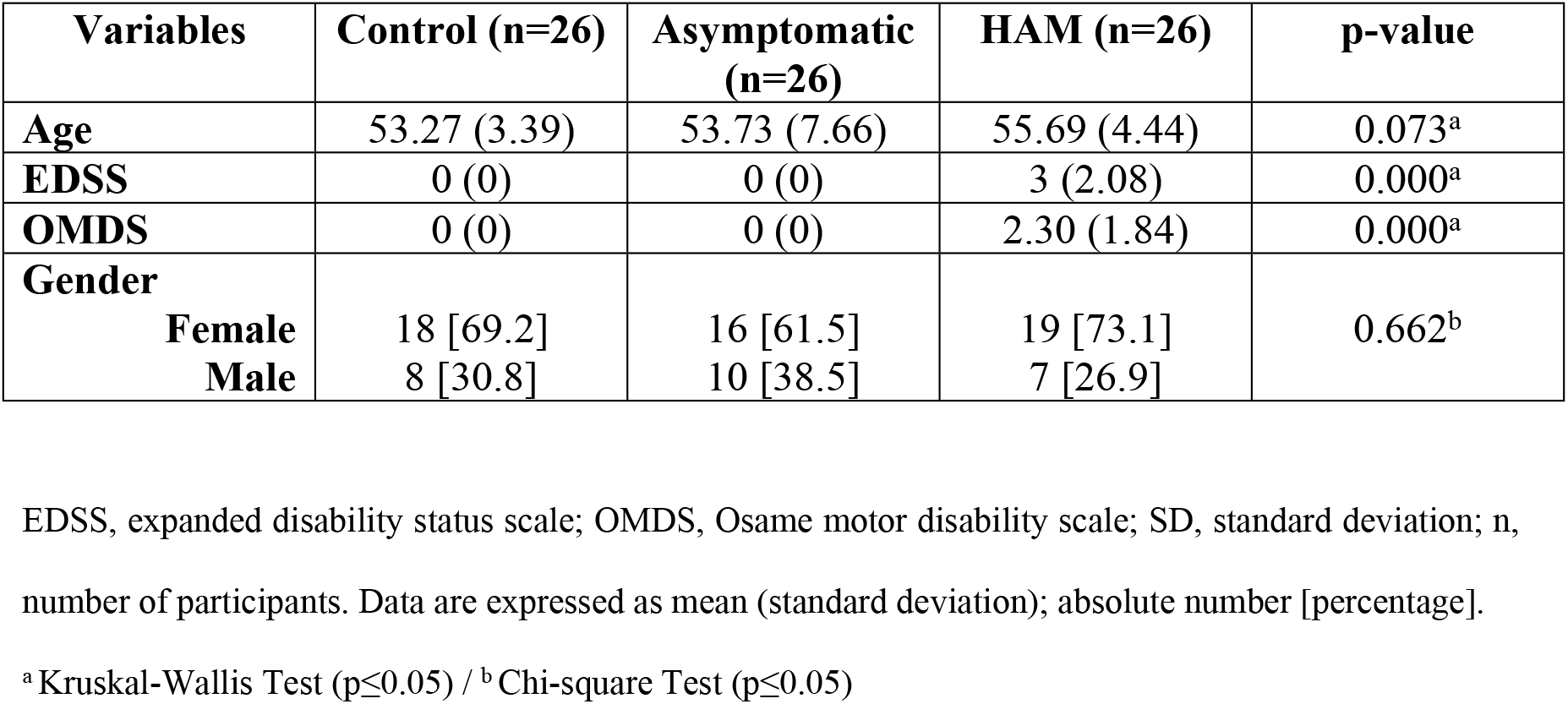
General characteristics and disability scales (EDSS and OMDS) of the participants (n=78)

| Variables | Control (n=26) | Asymptomatic (n=26) | HAM (n=26) | p-value |
| --- | --- | --- | --- | --- |
| **Age** | 53.27 (3.39) | 53.73 (7.66) | 55.69 (4.44) | 0.073<sup>a</sup> |
| **EDSS** | 0 (0) | 0 (0) | 3 (2.08) | 0.000<sup>a</sup> |
| **OMDS** | 0 (0) | 0 (0) | 2.30 (1.84) | 0.000<sup>a</sup> |
| **Gender** |  |  |  |  |
| **Female** | 18 [69.2] | 16 [61.5] | 19 [73.1] | 0.662<sup>b</sup> |
| **Male** | 8 [30.8] | 10 [38.5] | 7 [26.9] |  |

EDSS, expanded disability status scale; OMDS, Osame motor disability scale; SD, standard deviation; n, number of participants. Data are expressed as mean (standard deviation); absolute number [percentage].
<sup>a</sup> Kruskal-Wallis Test ( $p \leq 0.05$ ) / <sup>b</sup> Chi-square Test ( $p \leq 0.05$ )

The VEMP latencies were different among the groups. Table 2 indicates the comparative analysis and identifies the groups for which the difference was relevant.

**Table 2.** Comparison of the groups with HTLV-1-associated myelopathy, asymptomatic infection and healthy controls according to the latency (ms) of cervical VEMP and ocular VEMP. N=78

| Variables | G1<br>(n=26) | G2<br>(n=26<sup>a</sup>) | G3<br>(n=26<sup>b</sup>) | p -<br>value<br>* | Comparison<br>groups | p -<br>value ** |
| --- | --- | --- | --- | --- | --- | --- |
| Lat P13 | 12.8 (0.91) | 13.7<br>(1.03) | 14.8<br>(3.22) | 0.001 | G1 X G2<br>G1 X G3<br>G2 X G3 | 0.007<br>0.002<br>1.000 |
| Lat N23 | 22.3 (1.36) | 23.0<br>(2.44) | 25.7<br>(4.43) | 0.003 | G1 X G2<br>G1 X G3<br>G2 X G3 | 0.919<br>0.003<br>0.060 |
| Lat N10 | 10.5 (0.65) | 10.4<br>(0.92) | 11.5<br>(2.80) | 0.375 | - | - |
| Lat P15 | 15.4 (0.66) | 15.7<br>(1.35) | 18.2<br>(3.30) | 0.000 | G1 X G2<br>G1 X G3<br>G2 X G3 | 1.000<br>0.000<br>0.002 |

G1, group control; G2, group of asymptomatic individuals; G3, group of individuals with HAM; SD, standard deviation; Lat, latency; n, number of participants. Data are expressed as mean (standard deviation).
<sup>a</sup>For Lat N23 data analysis, one case in which the response was absent was excluded.
<sup>b</sup>For Lat P13 and Lat N23 data analysis, 3 and 6 cases, respectively, in which the response was absent were excluded. For Lat N10 and Lat P15 data analysis, 2 and 3 cases, respectively, in which the response was absent were excluded.
\* Kruskal-wallis Test ( $p \leq 0.05$ ) / \*\* Bonferroni Test

Table 3 describes the frequency of normal and altered results for cervical and ocular VEMP in each group.

**Table 3.** Comparison of cervical and ocular VEMP in the groups HTLV-1-associated myelopathy, asymptomatic infection and controls. N=78

| **Electrophysiological evaluation (VEMP)** | **G1 (n=26)** | **G2 (n=26)** | **G3 (n=26)** | **p - value \*** | **Comparison groups** | **p - value \*\*** |
| --- | --- | --- | --- | --- | --- | --- |
| Normal (cVEMP + oVEMP) | 26 [100] | 16 [61.5] | 0 [0] | 0.000 | G1 X G2<br>G1 X G3<br>G2 X G3 | 0.001<br>0.004<br>0.002 |
| cVEMP altered | 0 [0] | 7 [27] | 7 [27] | 0.689 | - | - |
| oVEMP altered | 0 [0] | 2 [7.7] | 3 [11.5] | 1.000 | - | - |
| cVEMP + oVEMP altered | 0 [0] | 1 [3.8] | 16 [61.5] | 0.000 | G1 X G2<br>G1 X G3<br>G2 X G3 | 0.060<br>0.003<br>0.004 |

G1, group control; G2, group of asymptomatic individuals; G3, group of individuals with HAM; cVEMP, cervical VEMP; oVEMP, ocular VEMP; n, number of participants. Data are expressed as absolute number [percentage].
\* Chi-square test or Fisher's Exact ( $p \leq 0.05$ ) / \*\* Bonferroni Test

The VEMP response was categorized as 1) latency delay of N10-P15 waves (ocular) or of P13-N23 waves (cervical); 2) absence of wave; 3) normal wave. Fig 3 shows the comparative analysis for ocular VEMP and Fig 4 for cervical VEMP.

**Fig 3.**
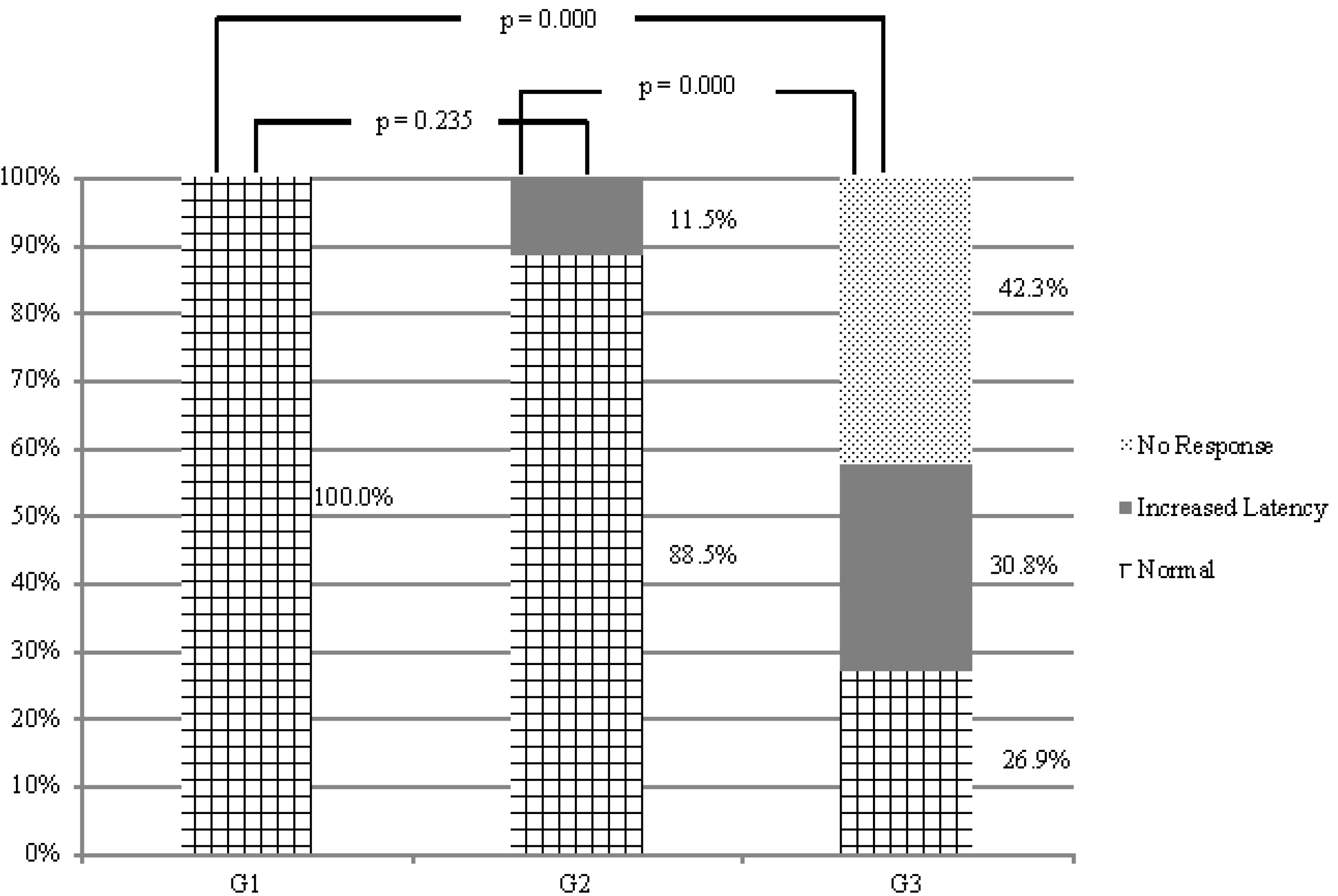
Comparison of ocular VEMP responses in individuals with HTLV-1-associated myelopathy, with asymptomatic infection and seronegative controls (n=78). G1, group control; G2, group of asymptomatic individuals; G3, group of individuals with HAM. Chi-square or Fisher’s Exact test (p≤0.05)

**Fig 4.**
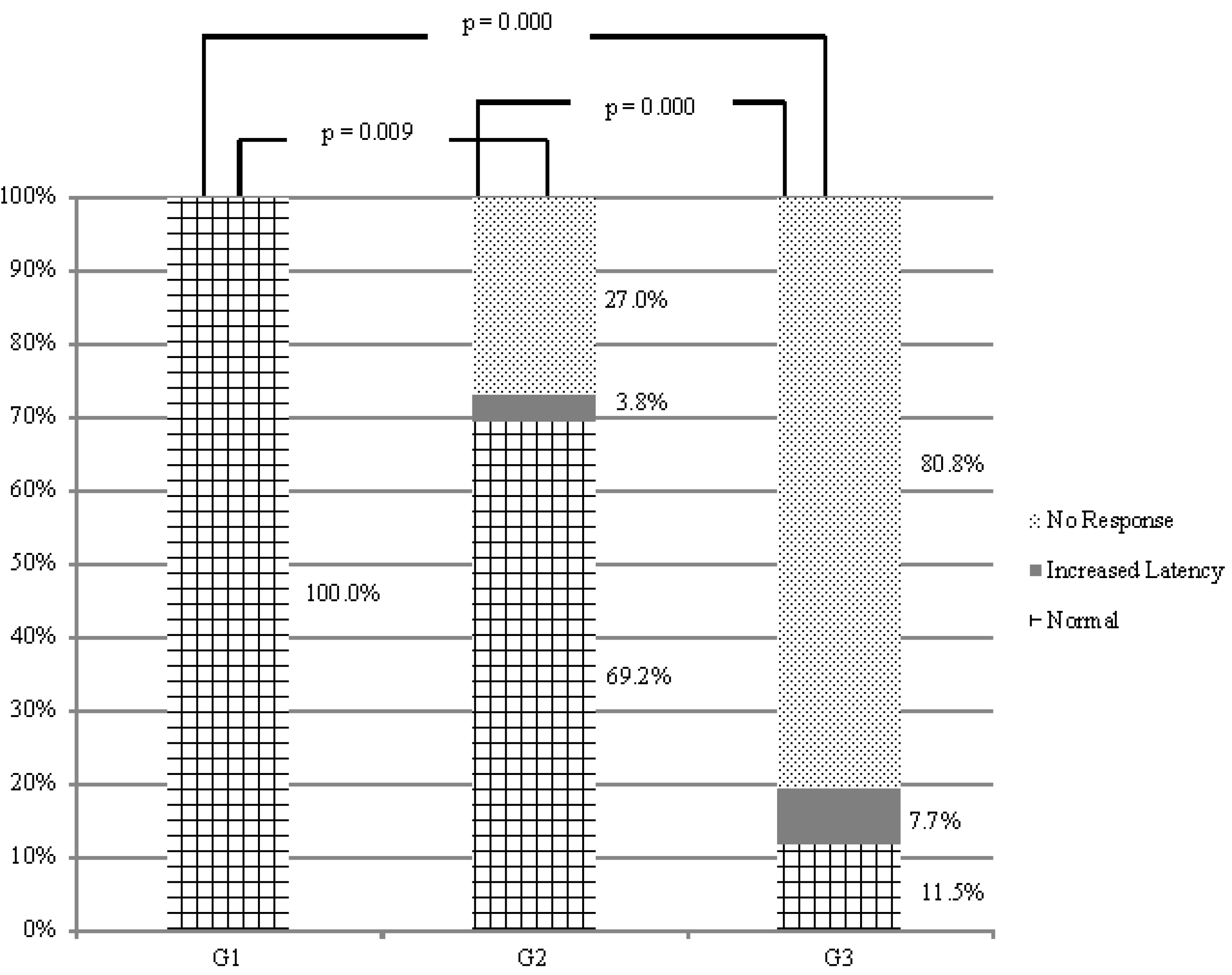
Comparison of cervical VEMP responses in individuals with HTLV-1-associated myelopathy, with asymptomatic infection and seronegative controls (n=78). G1, group control; G2, group of asymptomatic individuals; G3, group of individuals with HAM. Chi-square or Fisher’s Exact test (p≤0.05)

## Discussion

The auditory stimulus that evokes VEMP follows through the vestibular regions of the brain, especially the pre-motor cortex, the inferior and medial temporal gyrus, the Brodmann area, as well as the typically auditory areas, such as the primary auditory cortex [32].

The latency delay of cervical VEMP has been related to the demyelination of the primary afferent axon of the vestibulospinal tract and/or involvement of the vestibular nucleus [33–35]. The absence of electrophysiological response may be explained by a severe impairment of the vestibular-spinal pathway [36].

When the evoked potential changes from a prolonged latency to no response, it is understood that there is a worsening in the neuronal damage [14,27,28]. This pattern of response was previously observed in a cohort study of individuals infected by HTLV-1 with myelopathy and asymptomatic carriers that were tested by cervical VEMP [11].

Regarding cervical VEMP in the present study, we found that the great majority of the patients with definite HAM presented alteration in cervical VEMP response (88,5%). This data confirms previous studies that disclosed a cervical spinal cord damage in HAM, emphasizing that the medullary abnormalities in HAM are not restricted to the thoracolumbar level [37,38].

Regarding ocular VEMP, 61.5% of the patients with definite HAM and alteration in cervical VEMP, presented also alteration in ocular VEMP (Table 3). The neural connections involved in ocular VEMP are assumed to be thalamic and mesencephalic [22,39–41]. The presumed pathway includes the vestibular primary afferent, the vestibular nuclear complex, the medial longitudinal fasciculus, the oculomotor nucleus and the oculomotor nerves [39]. Thus, a latency delay or an absence of response depends on the disorganization of the primary afferents involved in the vestibulo-ocular reflex [39,40].

The higher frequency of simultaneous alteration in ocular and cervical VEMP of HAM group confirms a greater neurological impairment in these individuals with more advanced spinal cord injury when compared to the group with asymptomatic infection, although some individuals labeled as asymptomatic carriers were disclosed with altered VEMP. The meaning of this finding has to be studied as a possible signal predictive of HAM.

VEMP is able to detect subclinical neurological changes in HTLV-1 infection [16,19,32]. When effective therapeutic options for the HTLV-1 neurological disease are available, the subclinical diagnosis of neuronal injury will have implications in decision-making regarding the beginning of the treatment in the stage of incipient damage. For example, recent studies have shown that low doses of corticosteroid can be beneficial in slowing HAM progression if treatment is implemented at the onset of the HTLV-1-neurological manifestation [42,43].

## Conclusion

Ocular VEMP demonstrated that a subcortical impairment occurs in HAM and may occur even in the asymptomatic phase of HTLV-1 infection. Thus, neurological impairment in HAM is not restricted to the spinal cord. The vestibulo-ocular tract is subject to injury, compromising the oculomotor system in eye stabilization during head and body movements which can explain the high frequency of dizziness in patients with HAM.

## Supporting information

**S1 Table. Diagnostic criteria of human T-cell lymphotropic virus type 1 (HTLV-1)-associated myelopathy (HAM)^a^.** ^a^Castro-costa CMDE, Araújo AQC, Barreto MM, Takayanagui OM, Sohler MP, Silva ELMDA, et al. Proposal for diagnostic criteria of tropical spastic paraparesis/HTLV-1-associated myelopathy (HAM/TSP). AIDS Res Hum Retroviruses. 2006;22:931–935. Doi: 10.1089/aid.2006.22.931.

**S1 Fig. Questionnaire**

## Acknowledgments

We wish to thank the Interdisciplinary HTLV Research Group (GIPH) for the support.

